# Heterogeneity versus the COVID-19 Pandemic

**DOI:** 10.1101/2021.01.06.425543

**Authors:** Ramalingam Shanmugam, Gerald Ledlow, Karan P. Singh

## Abstract

In this paper, heterogeneity is formally defined, and its properties are explored. We define and distinguish observable versus non-observable heterogeneity. It is proposed that *heterogeneity* among the vulnerable is a significant factor in the contagion impact of COVID-19, as demonstrated with incidence rates on a Diamond Princess Cruise ship in February 2020. Given the nature of the disease, its heterogeneity and human social norms, pre-voyage and post-voyage quick testing procedures may become the new standard for cruise ship passengers and crew. The technological advances in testing available today would facilitate more humanistic treatment as compared to more archaic quarantine and isolation practices for all onboard ship. With quick testing, identification of those infected and thus not allowed to embark on a cruise or quarantining those disembarking and other mitigation strategies, the popular cruise adventure could be available safely again. Whatever the procedures implemented, the methodological purpose of this study should add valuable insight in the modeling of disease and specifically, the COVID-19 virus.

## 1. INTRODUCTION

In the literature, the term *heterogeneity* echoes differently in various contexts. What is heterogeneity or its antonym, homogeneity? Its root word lies in Greek “*heterogenes*” meaning different. In epidemiology or statistics disciplines, the word heterogeneity is popularly commented to exist when the variance is large. In insurance applications, for an example, the premium is assessed more if the insurer is in a heterogeneous group with high hazard proneness (Spreeuw, 1999). Should a large (small) variance be indicative of heterogeneity (homogeneity)? Interesting discussions are given for heterogeneity in Ecochard (2006); in healthcare disciplines, heterogeneity is referred to as different outcomes among patients. Should the heterogeneity be connected to only a non-observable hidden trait as done in genetics? Does heterogeneity refer dissimilar attributes across the subgroups of the population itself even before sampling? Is heterogeneity really pointing to the non-identical nature in a random sample or population? Should heterogeneity imply a shifting entity? In genetic studies, several authors refer to genetic heterogeneity as rather too difficult to ascertain. What do they really mean? If alleles in more than one locus exhibit susceptibility to a disease, there is a need to track the loci to infer their heterogeneity. So, in a sense, the application of heterogeneity is really a discussion of an opposite of similarity across loci. The reader is referred to Elston et al. (2003, pages 3404-344) for details. Hope and Norris (2013) attempted to determine how heterogeneity played a role in judgements in the context of crime victimization. Hence, what really is heterogeneity? A formal definition of heterogeneity is constructed later in the article, then, its properties are explored and itemized.

However, in the epidemiology literature, using a random sample *y*_1_, *y*_2_,…, *y*_*n*_ from a population whose main parameter is*θ*, when the null hypothesis *H*_*o*_: *θ*_1_ =*θ*_2_ = =*θ*_*n*_ is tested, it is named the *homogeneity test*. This suggests that heterogeneity is really all about a shifting population. This creates more confusion. Is the source of such confusion with respect to heterogeneity its ill communication? It is evident that there is a lack of a clear definition of heterogeneity given by Hunink et al. (2018, Chapter 12) for details. Neither the *Encyclopedia of Statistical Sciences n*or the *Encyclopedia of Biostatistics* has even an entry, as if it is not pertinent in statistical disciplines.

One comes across different types of data in epidemiologic studies. Drawing data from a binomial population is one of them, and the data should possess an under dispersion (i.e., variance of the binomial distribution is smaller than its mean). From a Poisson population, the drawn random sample ought to reflect equality between the mean and variance. When the main (incidence rate) parameter of a Poisson chance mechanism is stochastically transient, the unconditional observation of the random variable convolutes to an inverse binomial model (Ross, 2002). The inverse binomial distribution is known to attest that the variance is larger than its mean (Stuart and Ord, 2015, for details). Consequently, a comparison between the mean and variance characterizes only which type binomial, Poisson, or inverse binomial possesses the underlying chance mechanism we are sampling from but does not inform anything about heterogeneity.

With details about the probabilistic patterns among coronavirus confirmed, recovered, or cured individuals and those that succumb as fatalities/deaths in the thirty-two states/territories of India are given by Shanmugam (2020). To track the confusion with respect to heterogeneity, let us consider the data given in Table 1 (Mizumoto and Chowell, 2020), describing the spread of COVID-19 among the voyagers in a Diamond Princess Cruise ship, during the month of February 2020. The random variables *Y*_1_, *Y*_2_, and *Y*_3_ denote, respectively, the number of COVID-19 cases, the number of asymptomatic cases and the number of symptomatic cases among them in time (date). Under a given COVID-19’s prevalence rate,*λ* > 0, the number *Y*_1_ perhaps follows a Poisson probability pattern. For a given number of COVID-19 cases in a date, the number*Y*_2_ perhaps follows a binomial probability pattern with parameters (*y*_1_, *p*), where 0< *p* <1 denotes the chance for a COVID-19 case to exhibit no symptoms. Naturally, the number *Y*_3_ should follow a binomial probability pattern with parameters (*y*_1_,1− *p*). There is an implicitness between *Y*_2_ and *Y*_3_, in the sense that *Y*_2_ + *Y*_3_ = *Y*_1_. There are three-time oriented groups of COVID-19 incidences in Table 1. Is there an observable heterogeneity among the three groups? If so, is it due to a non-observable (parametric) heterogeneity? How do we define and distinguish observable versus non-observable heterogeneity? A literature search in epidemiology and/or biostatistics does not provide an answer to this question.

**Table 1.** COVID-19 in Cruise Ship, 2020, Mizumoto et al. (2020)

| Date | $Y_3$ | $Y_2$ | $Y_1$ | $\bar{\lambda} = \bar{y}_1$ | $s_{\lambda}^2 = \hat{V}ar(Y_1)$ | $OR_{\frac{1 \rightarrow 2}{0 \rightarrow 1}}$ | $Odds_{Y_1}$ |
| --- | --- | --- | --- | --- | --- | --- | --- |
| Feb 15-16, 2020 | 29, 32 | 38, 38 | 67, 70 | 68.5 | 4.5 | 0.5001 | 1.7E-30 |
| Feb 17-18, 2020 | 29, 23 | 70, 65 | 99, 88 | 93.5 | 60.5 | 0.5000 | 2.4E-41 |
| Feb 19-20, 2020 | 11, 7 | 68, 6 | 79, 13 | 46 | 21.78 | 0.5002 | 1.0E-20 |

It is evident that the average of COVID-19 cases is an estimate of COVID-19’s prevalence rate (i.e., 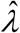 in Table 1). Their estimates impress that the prevalence rate is transient, not constant across every pair of two-day duration dyads. The Poisson population from which the COVID-19 cases are drawn ought to have been dynamic, implying the existence of a Poisson heterogeneity. How do we define and/or capture the heterogeneity level? This is the theme and purpose in this research article.

Likewise, given that a fixed number, *y*_1_ of COVID-19 cases has occurred, a part of them might be asymptomatic cases, *y*_2_ and the remaining are symptomatic cases, *y*_3_. That is, *y*_2_ and *y*_3_ are complementary but *y*_2_ + *y*_3_ = *y*_1_. Is there heterogeneity in each of the two sub-binomial populations, whether there is a heterogeneity in *y*_1_ ? How should each *binomial heterogeneity* be defined and computed? In other words, is binomial heterogeneity different from that of Poisson heterogeneity? If so, what are the differences? A literature search in epidemiology and/or biostatistics offers no help to prove either the existence or absence of binomial heterogeneity in the data for *y*_2_ or *y*_3_ in Table 1. Hence, we continue probing matters with respect to heterogeneity.

The concept of heterogeneity seems to have escaped the researchers and epidemiologists’ scrutiny for a long time. It is time well spent and worthwhile to revive an interest in the construct of heterogeneity, and that is exactly what this article is trying to accomplish. Hence, we first define and construct an approach for the idea of heterogeneity. To be specific, we first discuss Poisson heterogeneity and then take up binomial heterogeneity. Maybe our research direction about heterogeneity is, perhaps, pioneering. However, we believe that our approach is easily extendable for many other similar methodological setups. We illustrate our definition and all derived expressions for heterogeneity using COVID-19’s data pertaining to the Diamond Princess Cruise ship, Yokohama, 2020 as displayed in Table 1.

## 2. POISSON AND BIONOMIAL HETEROGENEITIES

Applied epidemiologists emphasize that heterogeneity is of paramount importance in extracting and interpreting data evidence. Many data analysts are convinced that an unrecognized heterogeneity leads to a biased inference. To begin with, what is heterogeneity? It is a factor causing non-similarities. If so, how many sources are there? We contemplate that there are two sources for heterogeneity to exist. One source ought to be from the drawn random sample of observations: *y*_1_, *y*_2_,….., *y*_*n*_, which we recognize as *observable heterogeneity*. Would the sampling variability, *Var*[*f* (*y*_1_, *y*_2_ ,….., *y*_*n*_ | *θ*)] for a selected statistic *f* (*y*_1_, *y*_2_ ,….., *y*_*n*_ *θ*) express the observable heterogeneity? Another source is manifested in non-observable parameter, *θ* of the chance mechanism, which we recognize as *non-observable heterogeneity*. Would a non-uniform stochastic pattern of*θ* be indicative of the non-observable homogeneity? If the chance mechanism perversely selects a probability density function (pdf) for*θ*, how would it manifest itself to portray the non-observable heterogeneity? Both observable and non-observable heterogeneity together ought to be involved to make any definition of heterogeneity complete. If so, how do we integrate them? Often, under/over-dispersion is confused with heterogeneity. It seems that the over/under dispersion is precipitated by heterogeneity but not the other way. It is not obvious or proven so far in the epidemiology literature on whether the converse is true. We focus only on Poisson and binomial populations to address heterogeneity, and these arguments can be repeated for other populations considering similar methods.

### 2.1. POISSON HETEROGENEITY

Recall that the random integer, *Y*_1_ denoting the number of COVID-19 cases in a place (like the Diamond Princess cruise ship) at a time (like February, 2020) is a Poisson random variable with a specified prevalence rate,*λ* > 0. That is, the conditional probability of observing *y*_1_ number of COVID-19 cases under a prevalence rate *λ* > 0 is 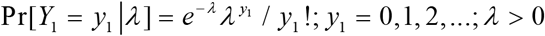 with its expected number *E* [*Y*_1_ |*λ*] = *λ* and variability*Var*[*Y*_1_ |*λ*] = *E*[*Y*_1_ ∣*λ*]. The reader is referred to Rajan and Shanmugam (2020) for detailed derivations of the Poisson mean and variance. The prevalence parameter *λ* itself is crucial in our discussions. The Poisson variability cannot be heterogeneity, because the expected value also changes when the variability changes due to their inter-relatedness. Realize that no two individuals on the ship are assumed to have the same level of susceptibility to the COVID-19 virus. It is reasonable to imagine that the prevalence levels follow a conjugate, stochastic gamma distribution. The so-called conjugate prior knowledge in the Bayesian framework smooths the statistical analytic process. It is known that the conjugate prior for the Poisson distribution is gamma, whose pdf is

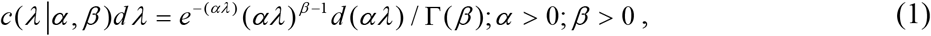

with an average. 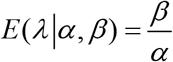 and variability *Var* (*λ∣α, β*) = *E* (*λ*∣*α, β*) / *α*, where the parameters*α*and *β* are recognized as *hyper-parameters* (Rajan and Shanmugam, 2020). Notice that the hyper parameter *α* > 0 causes the variability in the COVID-19’s prevalence rate to fluctuate up or down, and, hence, you would anticipate the heterogeneity to involve the hyperparameter*α*. But the question is how?

We assume that the probability of observing a non-negative COVID-19 case, *y*_1_ is a Poisson under a stable sampling population Pr(*Y*_1_ |*λ*) with an expected number *E* (*Y*_1_ ∣*λ*) = *λ*and a variability *Var* (*Y*_1_ ∣*λ*) = *E* (*Y*_1_∣*λ*). With replications, the observable heterogeneity should become estimable. That is to mention, the maximum likelihood estimate (MLE) of the COVID-19 prevalence rate is the average number, 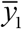, of the observations. To discuss the non-observable heterogeneity, we need to integrate its conjugate prior *c*(*λ*∣*α, β*) for the non-observable *λ* with the likelihood Pr(*Y*_1_ ∣*λ*) and it results in an update and it is called posterior pdf for*λ*. The expressions for non-observable heterogeneity, observable heterogeneity and other expressions are given in Appendix I.

### 2.2. BINOMIAL HETEROGENEITY

In this section, we explore heterogeneity for two sub-binomial processes emanating from a Poisson process. The asymptomatic number, *Y*_2_ and symptomatic number, *Y*_3_ of COVID-19 cases are two branching binomial random numbers out of the Poisson random number, *Y*_1_ = 0,1, 2,…; of COVID-19 cases. These two split random variables are complementary of each other in the sense that *Y*_2_ + *Y*_3_ = *Y*_1_. Then, what are the underlying model for*Y*_2_ and for*Y*_3_ ? Are they correlated random variables? If so, what is their correlation? These are pursued in this section.

Let *I* be an indicator random variable defined as: *I*_*i*_ = 1 for a COVID-19 case to be asymptomatic with a probability, 0< *p* <1 and *I*_*i*_ = 0 for the case to be symptomatic with a probability, 0 <1− *p* <1. Then, for a fixed *y*_1_, the random variable, 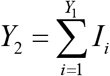 follows a binomial probability distribution with parameters (*y*_1_, *p*). Likewise, for a fixed *y*_1_, the random variable,*Y*_3_ = *y*_1_ − *Y*_2_ follows a complementary binomial distribution with parameters (*y*_1_,1− *p*) .That is,

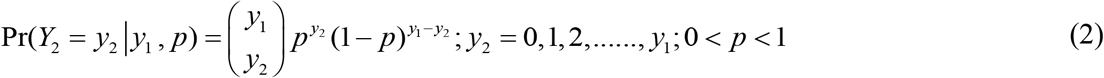

and

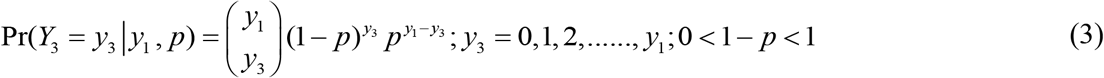

The expressions for non-observable heterogeneity, observable heterogeneity and other expressions are given in Appendix II.

## 3. TANGO INDEX

Lastly, we develop the *Tango index* and its significance level over the time period. Tango (1984) proposed an index to detect disease clusters in grouped data. This index received considerable attention in the literature. Following the line of thinking in Tango (1984), we could next assess the MLEs of several entities we estimated and displayed in Tables 1, 2, and 3. There are three groups of duration. Group 1 consists of the 15^th^ and 16^th^ of February 2020. Group 2 includes data for 17^th^ and 18^th^ of February 2020. Group 3 contains data of 19^th^ and 20^th^ of February 2020. Two independent contrasts among the three groups are feasible. In an arbitrary style, we select to compare Group 1 with Group 2 and then Group 2 with Group 3. For this purpose, we formulate a contrast matrix

**Table 2.** Results for Mizumoto et al.’s COVID-19 Data in Diamond Princess

| Date | $OR_{\frac{Y_3}{Y_2}}$ | $\hat{H}_{y_1}$ | $\hat{\beta}$ | $\hat{\alpha}$ | $d(y_1, \lambda)$ | $H_{\hat{\lambda}}$ |
| --- | --- | --- | --- | --- | --- | --- |
| 15, 16 Feb 2020 | 943.88 | **0.27** | 15.22 | 1042.72 | 857.81 | 0.93 |
| 17, 18 Feb 2020 | 7.36E+17 | **0.70** | 1.54 | 144.50 | 18.79 | 0.61 |
| 19, 20 Feb 2020 | 9.69E+11 | **0.65** | 2.11 | 97.15 | 23.56 | 0.67 |

**Table 3.** Results for Asymptomatic COVID-19 Cases in Mizumoto et al. (2020)

| Date | $1 - \bar{p} =$<br>$1 - Ave(\frac{y_2}{y_1})$ | $s_p^2$<br>$= \hat{V}ar(\frac{y_2}{y_1})$ | $\hat{H}_{y_1, \gamma, \delta}$ | $\hat{H}_{y_2\vert y_1}$ | $d(y_2, p)$ | $d(Y_2, Y_3)$ |
| --- | --- | --- | --- | --- | --- | --- |
| 15, 16 Feb 2020 | 0.45 | 0.0002 | **0.99** | **0.95** | 37.125 | 6.85 |
| 17, 18 Feb 2020 | 0.28 | 0.0004 | **0.98** | **0.89** | 66.6 | 41.14 |
| 19, 20 Feb 2020 | 0.34 | 0.0796 | 0.10 | **0.74** | 29.7 | 14.72 |

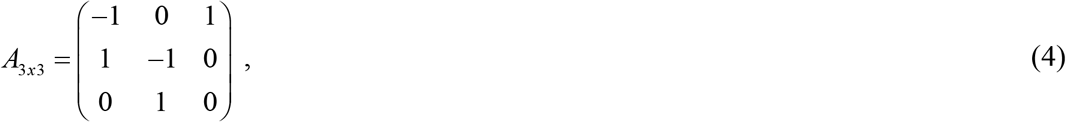

where the third column of the matrix needs no explanation. The Tango’s statistic*T* = *<u>r</u>* ’ *A<u>r</u>* follows a chi-square distribution with *v* = 2 degrees of freedom (df), where 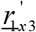 is a row vector of the MLE of a chosen entity in our analytic results in Table 1 or Table 2 or Table 3. For an example, let *<u>r</u>* ’ = (68.5,93.5,46) for the MLE of the COVID-19 prevalence rate,*λ* in the groups. Then, the Tango’s test statistic is *T* = 422.25 with *v* = 2 df and *p* − *value* = 2.03975*E* −92. Likewise, the Tango’s test statistic value and its p-value are calculated and displayed in Table 4 for other entities.

**Table 4.** Tango’s Test Statistic and Its P-Value for Several Entities

| Tango<br>statistic | $\hat{H}_{y_1}$ | $H_{\hat{\lambda}}$ | $\bar{p}$ | $\hat{H}_{y_1, \gamma, \delta}$ | $\hat{H}_{y_2\vert y_1}$ | $d(y_1, \lambda)$ | $d(y_2, p)$ | $d(Y_2, Y_3)$ |
| --- | --- | --- | --- | --- | --- | --- | --- | --- |
| T with 2 df | 0.25 | 0.36 | 0.41 | **0.77** | **0.51** | 699420 | 260.66 | 751.20 |
| p-value | 0.87 | 0.83 | 0.81 | **0.67** | **0.77** | 0.0E100 | 2.4E-57 | 7.5E-164 |

## 4. ILLUSTRATING USING COVID-19 DATA OF THE DIAMOND PRINCESS CRUISE SHIP

In this section we illustrate all the concepts and expressions of Section 2. Let us consider the COVID-19 data in Table 1 for the Diamond Princess Cruise Ship, 2020. The Diamond Princess is a cruise ship registered in Britain and operated across the globe. During a cruise that began on 20 January 2020, positive cases of COVID-19 linked to the pandemic were confirmed on the ship in February 2020. Over 700 people out of 3,711 became infected (567 out of 2,666 passengers and 145 out of 1,045 crew), and 14 passengers died. To be specific, on the 15^th^ of February 2020, 67 people were infected, on the 16^th^ of February 2020, 70 people were infected, on the 17^th^ of February 2020, there were 99 COVID-19 cases, on the 18^th^ of February, another 88 cases were confirmed. The U.S. government initially asked Japan to keep the passengers and crew members on board the ship for 14 days. The U.S. government, however, later decided to bring them to an Air Force base in California and a base in San Antonio, Texas.

For each specified day in the first column in Table 1, the estimate of COVID-19’s prevalence rate and its variance are calculated using expressions 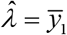 and 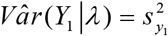. Both the prevalence and its variability increased and then decreased over the days. However, their correlation, 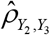 is calculated using the observed numbers on *y*_2_ and *y*_3_ for each day (see in Table 2) and the estimated correlations had been stable over the days. Substituting 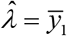 and 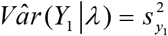 in the expression

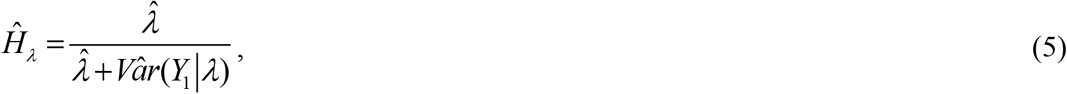

we obtained the non-observable heterogeneity and displayed in Table 2. The non-observable Poisson heterogeneity for *y*_1_ was high on the beginning day, came down later, and then increased. Using 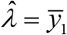 and 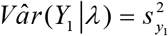 in the expression

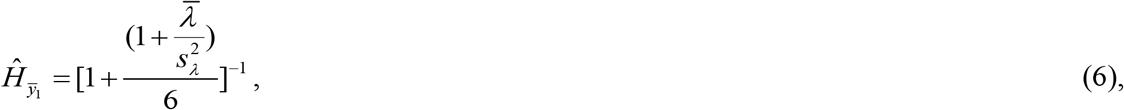

we obtained the observable heterogeneity and displayed in Table 2. The observable Poisson heterogeneity was low on the first day, increased and then decreased. Note in Table 2 that the observable and non-observable Poisson heterogeneities are inversely proportional. In other words, the estimate of the shape and scale parameter in the Bayesian approach are respectively 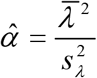 and 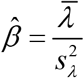 (see their values in Table 2). The shape parameter value decreased consistently over the days. The scale parameter was high to begin with, then increased later. The distance, *d* (*y*_1_,*λ*) between the observable and non-observable Poisson mechanism for *y*_1_ is calculated using the expression

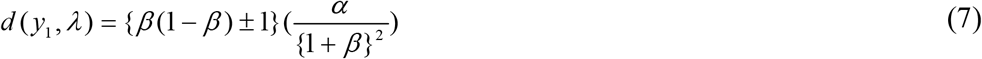

and displayed in Table 2. Notice that the distance was large to begin with, then decreased but increased later over the days.

Note that we compute 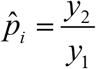 for the *i*^*th*^ day. Then, we calculate the

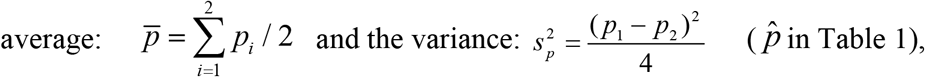

and it had been steadily increasing over the days since 15^th^ February 2020. This is something valuable for medical professionals learning the clinical nature of COVID-19. Using the expression,

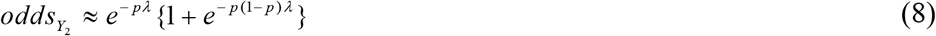

in Section 2.2, we calculated the odds for a COVID-19 case to become an asymptomatic type and displayed in Table 2.

Likewise, using the expression

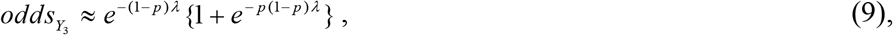

we estimated the odds for a COVID-19 case to become a symptomatic case as shown in Table 2. Notice that both odds (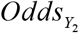 and 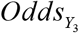) are low but their odds ratio,

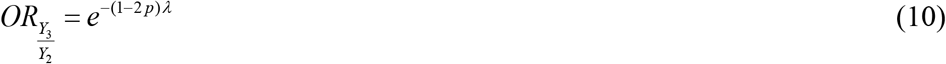

is not negligible but reveals that the situation is favorable to symptomatic rather than asymptomatic. This discovery is feasible because of the approach, and it is an eye-opening reality for the medical professionals in their desire to control the spread of the COVID-19 virus. Both the observable, 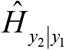 and non-observable, 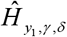 binomial heterogeneity (see their values in Table 3) were decreasing for the number, *y*_2_ of asymptomatic COVID-19 cases. The distance, *d* (*y*_2_, *p*) between the observable and non-observable for asymptomatic cases was moderate in the beginning, then increased, and then decreased over the next days (see their values in Table 3). However, the distance, *d* (*Y*_2_,*Y*_3_) between the observable, *y*_2_ of the asymptomatic cases and the observable, *y*_3_ of the symptomatic cases was narrow, then wider, and then moderate over the days (their values in Table 3).

For a COVID-19 case to become a symptomatic type, the chance is moderate to less and then more over the days (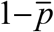 in Table 3). The estimate of the shape and scale parameter happened to be 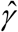and 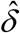 respectively (see their values in Table 3). Both the shape parameter and the scale parameter values decreased drastically over the days. From the p-values in Table 4, we infer that the prevalence rate, 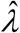, the distances, *d* (*y*_1_,*λ*), *d* (*y*_2_, *p*) and *d* (*Y*_2_ ,*Y*_3_) do differ significantly over the three groups of dyad days. The chance for COVID-19 to become an asymptomatic type does not differ significantly across the three groups. On the contrary, the non-observable heterogeneities 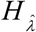 of the Poisson random number, *y*_1_ and 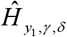 of the binomial random number, *y*_2_ are not significant. Likewise, the observable heterogeneities 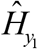 of the Poisson random number, *y*_1_ and 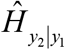 of the binomial random number, *y*_2_ for a given *y*_1_ are not significant.

## 5. DISCUSSION AND CONCLUSION

The risk of contracting the COVID-19 virus during a cruise is more than in a community setting, as confined spaces discourage non-pharmaceutical mitigation strategies such as social distancing to be weakly implemented and breathing air is tightly internalized. More nations are afraid to let the voyagers come ashore at the seaports. Ships are not even permitted to dock at the port, as to not complicate virus mitigation efforts by the local surrounding communities. The scenario seems to be anti-humanistic. The medical doctors and/or pharmaceutical service were strained due to the infected and COVID-19-free voyagers. Lack of clear symptoms among those that were infected added to difficulties in managing the COVID-19 crisis onboard the ship, and for any ship for that matter. Most importantly, how do we dispose of the COVID-19 fatalities (bodies), in a safe manner?

In the midst of uncertainties about the root cause and/or the appearance of any symptoms, the best modelers can do (as it is done in this article) is to devise a methodology to address the observable as well as non-observable heterogeneity, estimate the proportion of COVID-19 cases to be asymptomatic, estimate the odds of becoming symptomatic, and also the odds ratio for asymptomatic in comparison to those symptomatic among COVID-19 cases. Some of these are non-trivial to the professional experts dealing with the intention of reducing the spread of COVID-19 if not its total control. Still much of COVID-19 is a mysterious pandemic. It is clear that non-pharmaceutical mitigation strategies such as social distancing, utilization of face coverings, frequent hand sanitization, infected people quarantining on board, and severely controlled ship cleanliness and sanitation standards are required; this may only be successful with limited numbers of passenger and crew members. Given the nature of the disease, its heterogeneity and human social norms, pre-voyage and post-voyage quick testing procedures may become the new standard for cruise ship passengers and crew. The technological advances in testing provided today would facilitate more humanistic treatment as compared to more archaic quarantine and isolation practices for all onboard ship. With quick testing, identification of those infected and thus not allowed to embark on a cruise or quarantine those disembarking, and other mitigation strategies, the popular cruise adventure could be available safely again. Whatever the procedures implemented, the methodological purpose of this study should add valuable insight in the modeling of disease and specifically, the COVID-19 virus.

## Availability of data and materials

There is no other data or materials other than what are in the manuscript itself.

## Code availability

None.

## Authors’ contribution

The authors contributed everything in the manuscript.

## Acknowledgements

The authors thanks Texas State University and The University of Texas Health Science Center at Tyler for the support to write this article.

## APPENDIX I

### Poisson Heterogeneity: Derivations

It is known that the conjugate prior for the Poisson distribution is gamma, whose pdf is

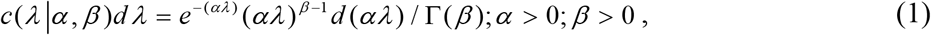

with an average. 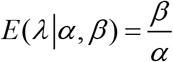 and variability *Var* (*λ*∣*α,β*) = *E* (*λ*∣*α,β*) / *α*, where the parameters*α* and *β* are recognized as *hyper-parameters* (Rajan and Shanmugam, 2020). Notice that the hyper parameter*α* > 0 causes the variability in the COVID-19’s prevalence rate to fluctuate up or down and hence, you would anticipate the heterogeneity to involve the hyperparameter*α*.

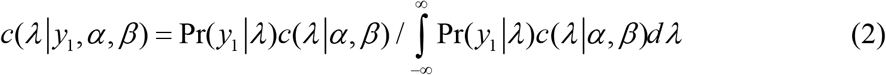

is the posterior pdf of the non-observable *λ*. Also, the denominator

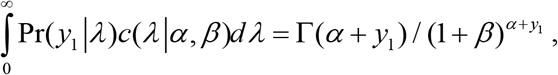

in a Bayesian framework, is called the *marginal distribution*. With Δ_*λ*_ = *λ*− *E*(*λ*), it is clear that 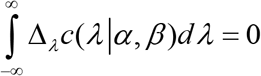, note that the prior variance is

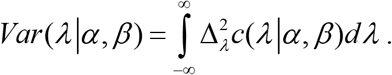

Because the prior is conjugate, its counterpart’s variability

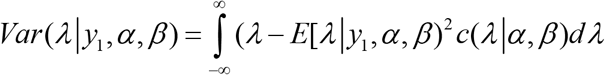

is minimal when the Bayes estimate of the non-observable is the posterior mean,*λ*_*Bayes*_ = *E*[*λ* ∣ *y*_1_*α, β*], where

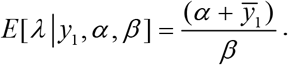

Differentiating the log-likelihood function

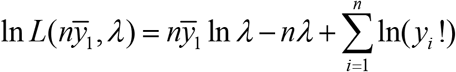

with respect to the non-observable parameter,*λ*, setting it equal to zero and solving it, we obtain the MLE and it is 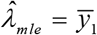. Because of the invariance property of the MLE, it is involved. The invariance property refers to that the MLE of a function of the parameter is the function of the MLE of the parameter. Also, it is known (Blumenfeld, 2010) that

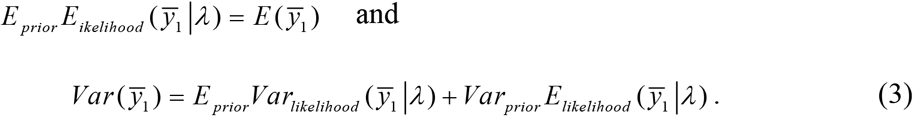

Hence, we are ready now to define the non-observable heterogeneity below in the Definition 1.

*Definition 1*. The non-observable heterogeneity of the Poisson parameter, *λ* is defined as

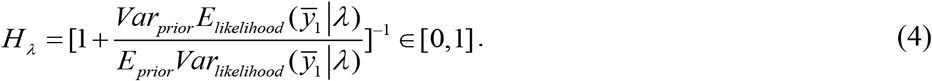

Following the Definition 1, we obtain the *non-observable heterogeneity* of the COVID-19 cases is

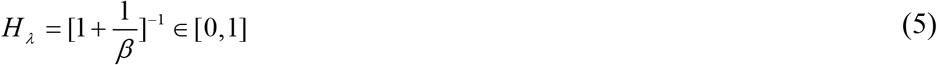

When the value of *H*_*λ*_is closer to zero, the data are believed to have non-observable Poisson homogeneity. Its MLE is

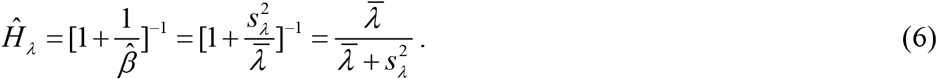

The reader is referred to Figure 1 for the configuration of the non-observable Poisson heterogeneity in general.

**Figure 1.**
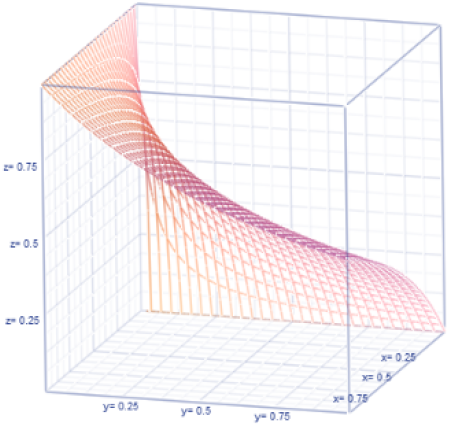
Non-observable Heterogeneity

Likewise, the *observable-heterogeneity* is defined below in Definition 2.

*Definition2*. The observable heterogeneity of the randomly sampled Poisson counts, *y*_1_, *y*_2_,….., *y*_*n*_ is defined as

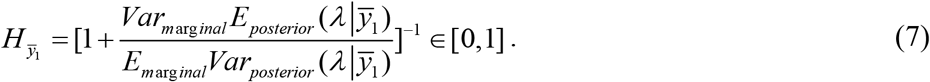

Before we apply the Definition 2, let us recollect that the marginal pdf of the complete sufficient statistic, 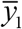 is uniform distribution and the posterior distribution is

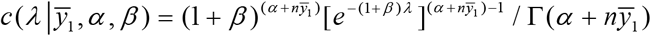

with

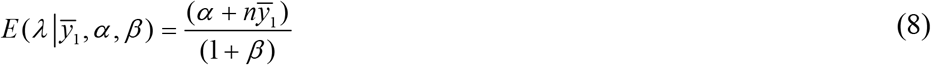

and

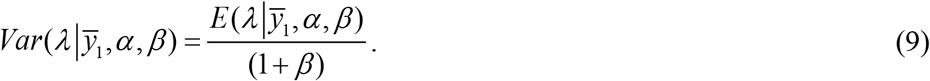

Imposing the Definition 2 and simplifying, we obtain that 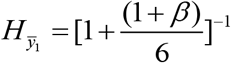 whose MLE is

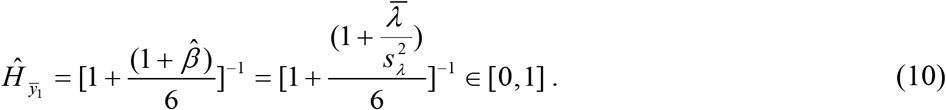

The reader is referred to Figure 2 for the configuration of the observable Poisson heterogeneity, 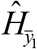 in general. When the value of 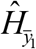 is closer to zero, the data are interpreted to have observable homogeneity.

**Figure 2.**
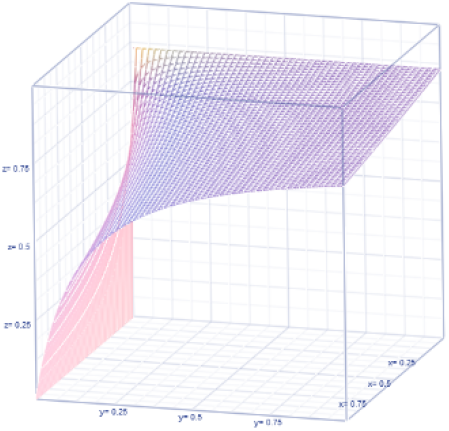
Observable Heterogeneity

Furthermore, the distance, *d* (*y*_1_,*λ*) between the observable *y*_1_ of the number of COVID-19 cases and the prevalence rate *λ* could be assessed using the formula

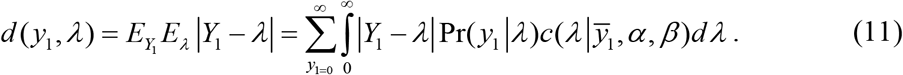

Realizing that their absolute difference is really ∣*Y*_1_ − *λ* ∣= *Y*_1_ + *λ*− 2 min{*Y*_1_,*λ*}, we obtain after simplifications that

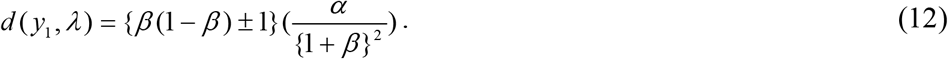

The configuration of the distance, *d* (*y*_1_,*λ*) between the observable and non-observable in Poisson mechanism. We now turn to discuss stochastic properties of the Poisson distribution are given in Figure 3.

**Figure 3.**
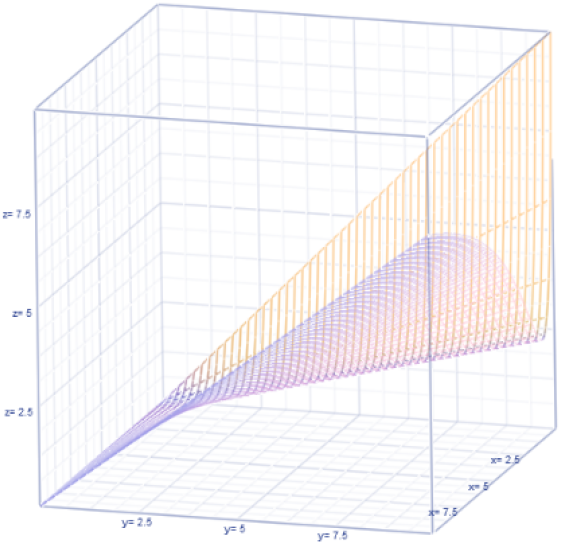
Distance, *d* (*y*_1_,*λ*) in Poisson.

The survival function of the random number, *Y*_1_ of COVID-19 cases is

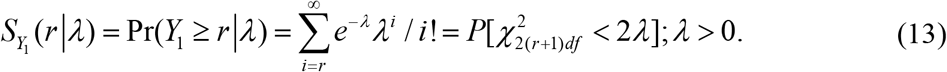

The hazard rate is a force of mortality. The hazard rate, *h*(*y*_1_) for the COVID-19 occurrence is

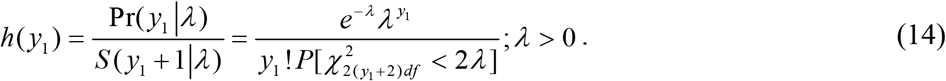

Does the Poisson chance mechanism keep any a finite *memory?* For example, the geometric distribution is known to have no memory. What is memory? The memory is really a conditional probability. That is,

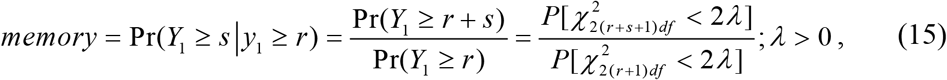

confirming that there is a finite memory in the Poisson mechanism of COVID-19 incidences. To be specific, with *r* = 0, *s* =1in the above result, the memory between COVID-19 free situation and just one COVID-19 occurrence is revealed in the chance-oriented Poisson mechanism. Such a memory is

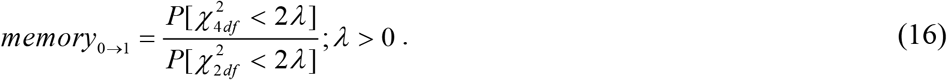

Likewise, the memory between at least one COVID-19 case situation and at least two COVID-19 cases situation is revealed with a substitution of *r* =1,*s* =1 in the above result and it is

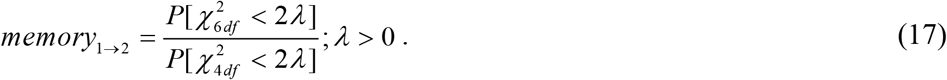

The odds ratio from the initial *memory*_0→1_ to the next *memory*_1→2_ is

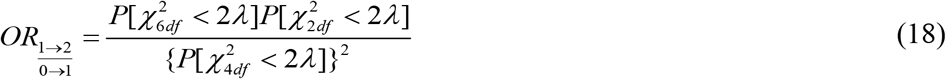

(their values in Table 1). However, the odds for COVID-19 free healthy situation to prevail is

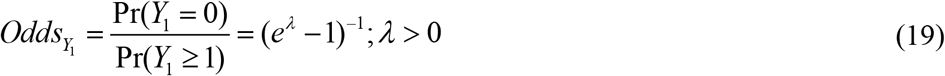

(their values in Table 1). For details on how the chance for an incidence of a disease to occur from a disease-free scenario changes, the reader is referred to Shanmugam and Radhakrishnan (2011).

## APPENDIX II

### Binomial Heterogeneity: Derivations

Let an indicator random variable, *I*_*i*_ = 1 for a COVID-19 case to be asymptomatic with a probability, 0< *p* <1and *I*_*i*_ = 0 for the case to be symptomatic with a probability, 0 <1− *p* <1. Then, for a fixed *y*_1_, the random variable, 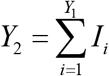 follows a binomial probability distribution with parameters (*y*_1_, *p*). Likewise, for a fixed *y*_1_, the random variable, *Y*_3_ = *y*_1_ − *Y*_2_ follows a complementary binomial distribution with parameters (*y*_1_,1− *p*) .That is,

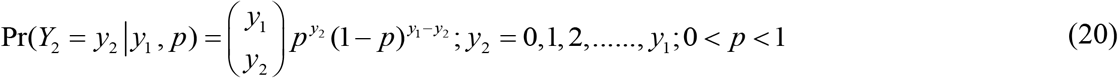

and

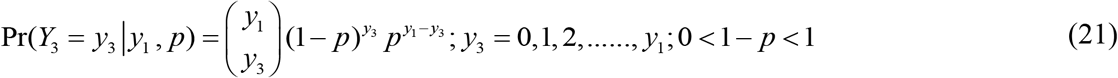

with their conditional expected numbers

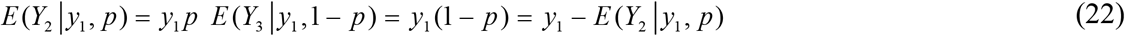

and the conditional variabilities

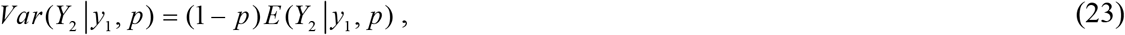

and

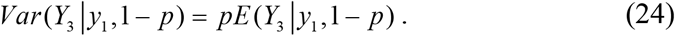

The conditional variability of*Y*_2_ is a percent (1− *p*) of its expected number *E* (*Y*_2_ ∣*y*_1_, *p*), implying that it exhibits under dispersion. Likewise, the conditional variability of*Y*_3_ is a percent (1− *p*) of its expected number *E* (*Y*_3_ ∣*y*_1_, *p*) = *y*_1_ (1 − *p*) implying that it also exhibits under dispersion. Together, the above statements suggest a conditional balance

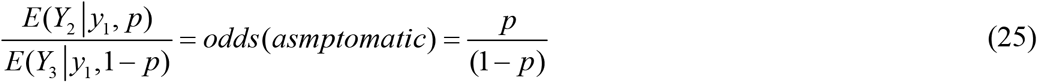

(Stuart and Ord, 2015 for details of the odds concepts). Consequently, we note that

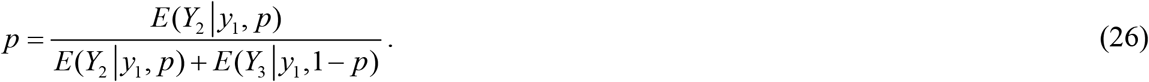

Furthermore, we wonder whether the random variables *Y*_2_ and *Y*_3_ are correlated? The answer is affirmative. To identify their correlation, notice that

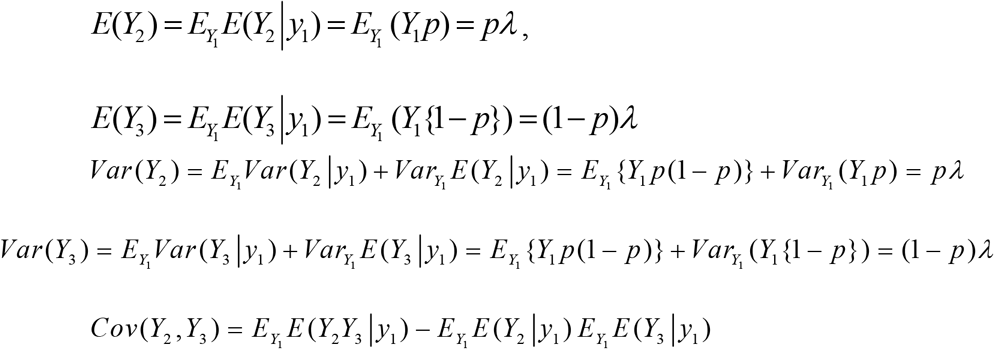

where

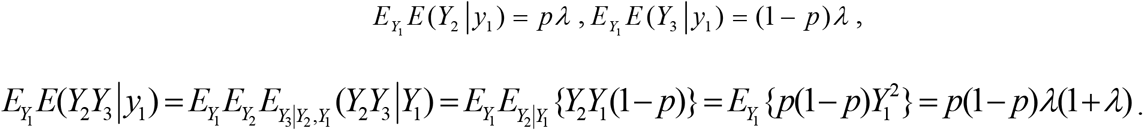

Hence, their correlation is

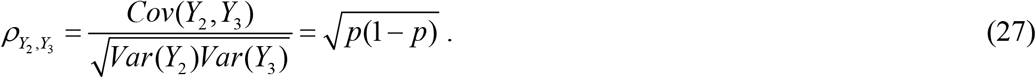

Their expected distance, 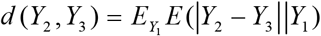 portrays the drift between the symptomatic observable,*Y*_2_ and the asymptomatic observable, *Y*_3_ and it is simplified to this function *d* (*Y*_2_, *Y*_3_) = ∣2 *p* − 1∣ *λ* (see Table 3 for their values), due to applying

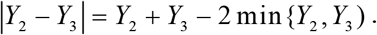

Let us assume that every COVID-19 case has the same chance of being asymptomatic in a time period. Then, the random number, *y*_2_ for a specified number, *y*_1_ of COVID-19 cases follows a binomial distribution with parameters (*y*_1_, *p*). We select a conjugate beta prior distribution

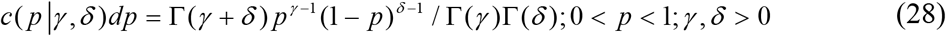

for our discussion for asymptomatic COVID-19 cases. The prior average is

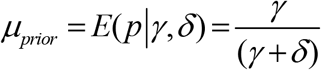

and the prior variability is

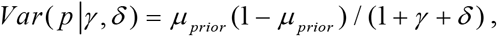

where the parameters*γ* and*δ* are hyper-parameters (Rajan and Shanmugam, 2020, for details). We guess that the binomial heterogeneity would involve both hyper parameters. The task for us is how do we construct such heterogeneity? An answer is the following. The posterior distribution

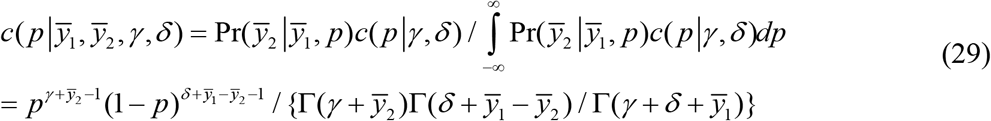

would play a key role to construct both the observable and non-observable binomial heterogeneity. With Δ _*p*_ = *p* − *E*(*p*), it is clear that 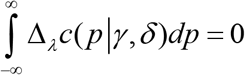.

The prior variance is

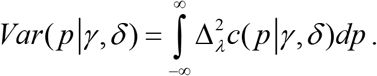

Its posterior counterpart

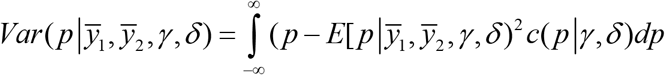

is minimal when the Bayes estimate of non-observable is the posterior mean

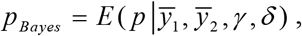

where

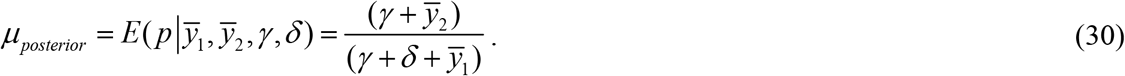

The posterior variance is

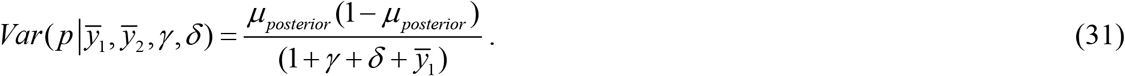

Differentiating the log-likelihood function as

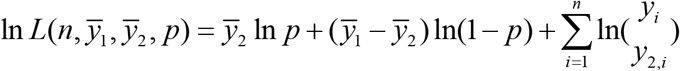

with respect to the non-observable parameter, *p*, setting it equal to zero and solving it, we obtain the MLE and it is 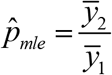 It is known that

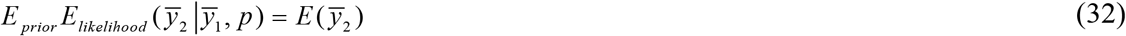

and

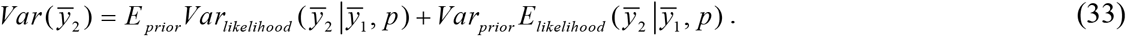

Hence, we define the non-observable binomial heterogeneity below in Definition 3.

*Definition 3*. The non-observable binomial heterogeneity is defined as

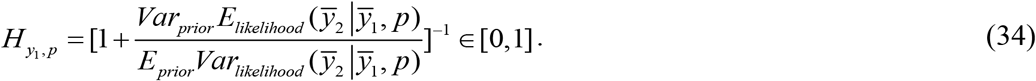

Following the Definition 3, we obtain the *non-observable heterogeneity* of the COVID-19’s asymptotic cases (remembering that (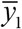,*γ,δ*) are the non-observable parameters) as

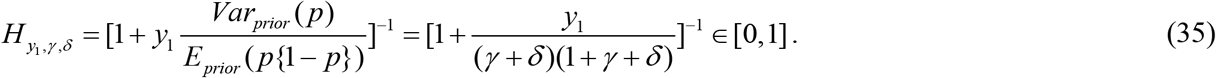

When the value of 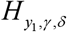 is closer to zero, the data are interpreted to have non-observable binomial homogeneity. Substituting the MLEs

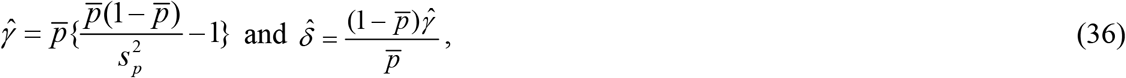

we obtain its MLE

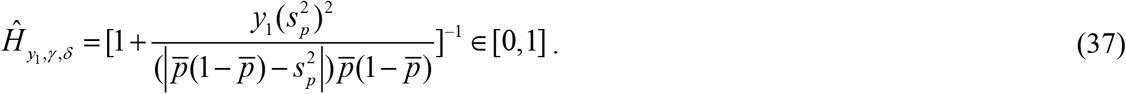

Likewise, the *observable-heterogeneity* of the binomial distribution of *y*_2_ is defined below in Definition 4.

*Definition 4*. The observable heterogeneity of the binomial counts, *y*_2,*i*_, *i* = 1, 2,…, *y*_1_ (in terms of the complete sufficient statistic 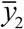) is defined as

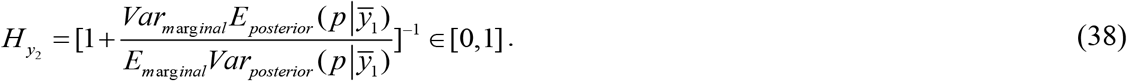

Before we apply Definition 4, remember that the marginal pdf of the complete sufficient statistic, 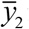 is the beta-binomial distribution,

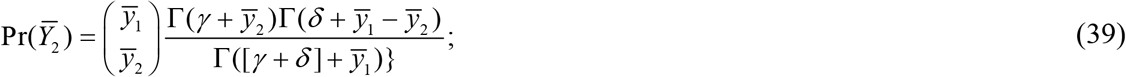

and the posterior distribution is beta. With the notation 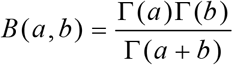, we note that the probability mass function of the beta-binomial distribution is

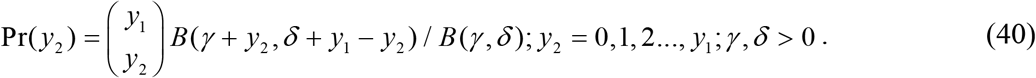

That is, the posterior probability density function is

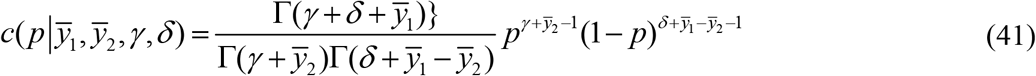

with

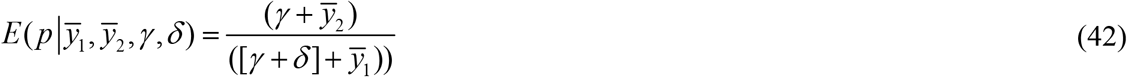

and

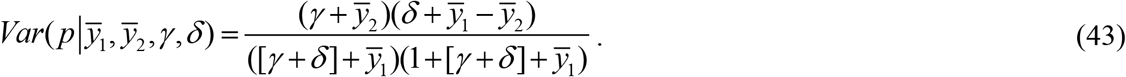

Now applying Definition 4, we obtain an expression for the observable binomial heterogeneity

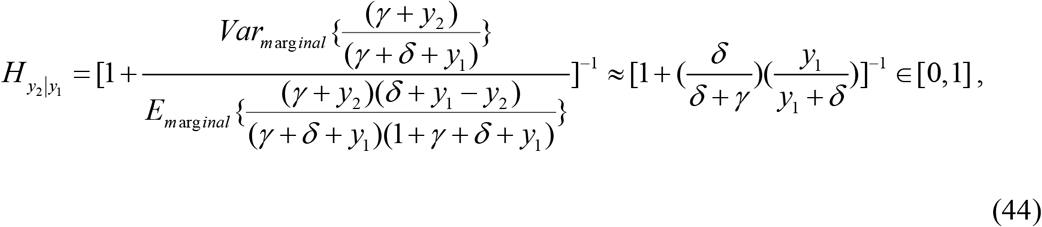

whose estimate is

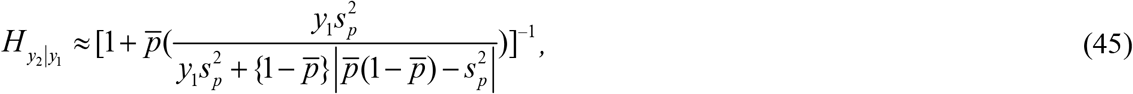

Because

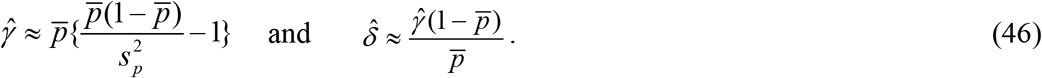

When the value of 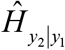 is closer to zero, the data are considered to have observable binomial homogeneity. Also, the distance, *d* (*y*_2_, *p*) between the observable *y*_2_ of the number of asymptomatic COVID-19 cases and its proportion, *p* could be assessed using the formula

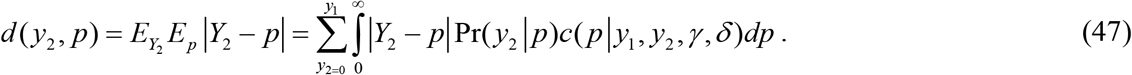

Realizing that the absolute difference, ∣*Y*_2_ − *p* ∣= *Y*_2_ + *p* − 2 min{*Y*_2_, *p*}, we obtain after simplifications that

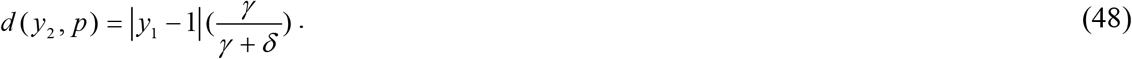

Likewise, to obtain the *non-observable heterogeneity* of the COVID-19’s symptomatic cases, all we have to do is change *p* to (1− *p*), change *y*_2_ to *y*_3_, along with changing*γ* to*δ* and go through the process above. Hence, the non-observable heterogeneity in the symptomatic cases is the same. That is,

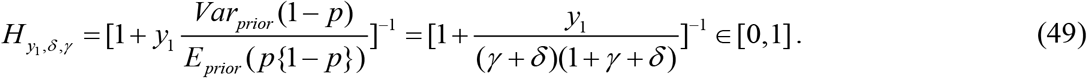

The observable binomial heterogeneity for the symptomatic cases is

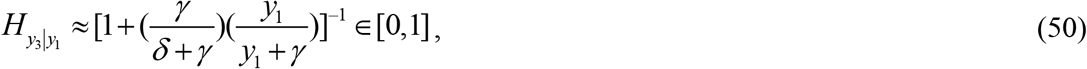

whose MLE is

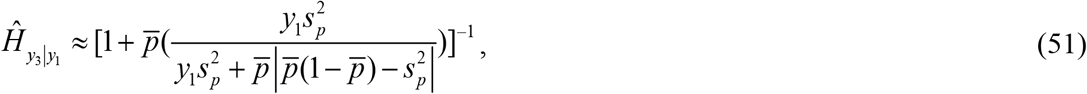

which is interestingly not the same as 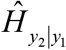. Also, the distance, *d*(*y*_3_,1− *p*) between the observable *y*_3_ of the number of asymptomatic COVID-19’s symptomatic cases and the proportion,1− *p* could be assessed using the formula

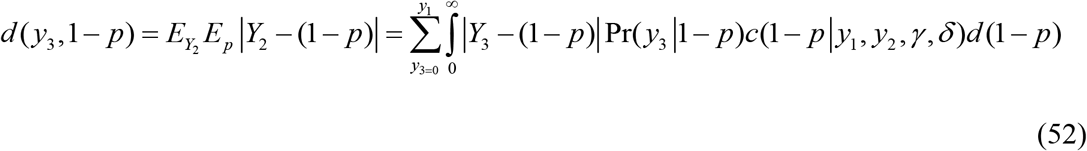

and it is after simplifications that

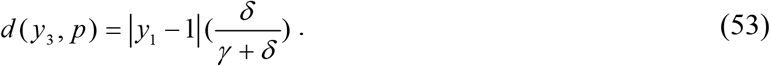

Now we explore statistical properties of the asymptomatic cases, *y*_2_. The survival function of the random number, *Y*_2_ with asymptotic symptoms is

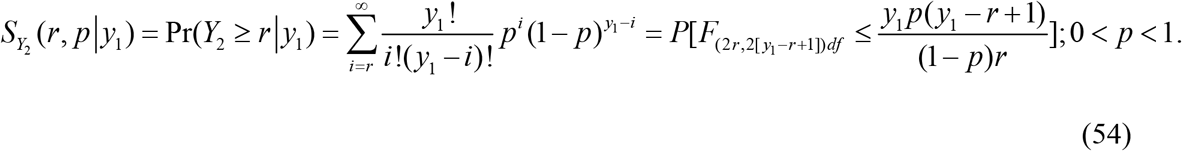

The hazard rate, *h*(*y*)of the binomial distribution for the asymptomatic cases is

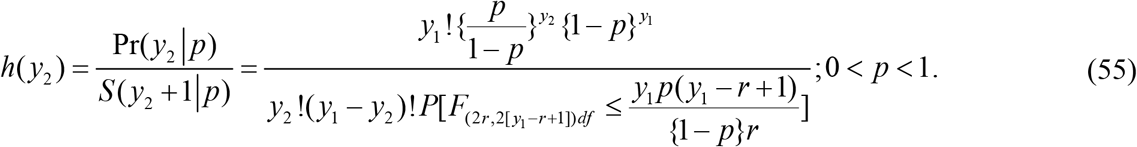

The binomial distribution has a finite *memory*

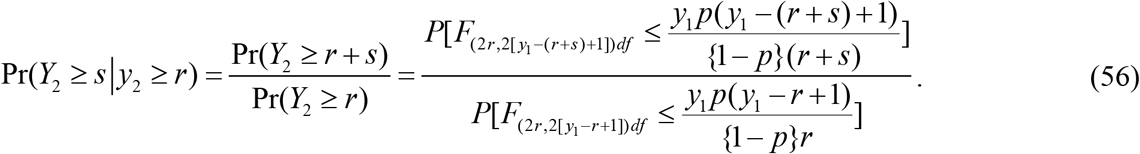

confirming that the usual binomial distribution does possess a finite memory. The conditional odds, for a fixed *y*_1_, for *safe* asymptomatic symptom are

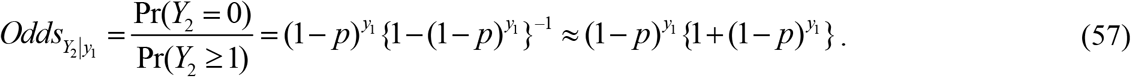

The unconditional odds for safe asymptotic symptom are

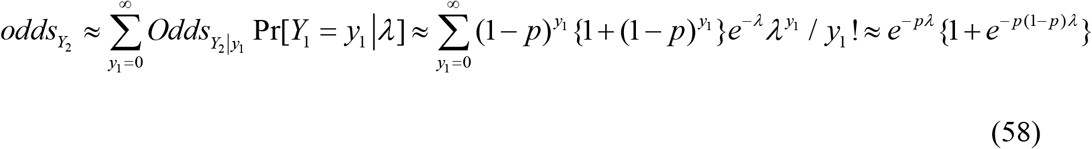

The reader is referred to Figure 4 for the configuration of the odds in asymptotic COVID-19 occurrences in general.

**Figure 4.**
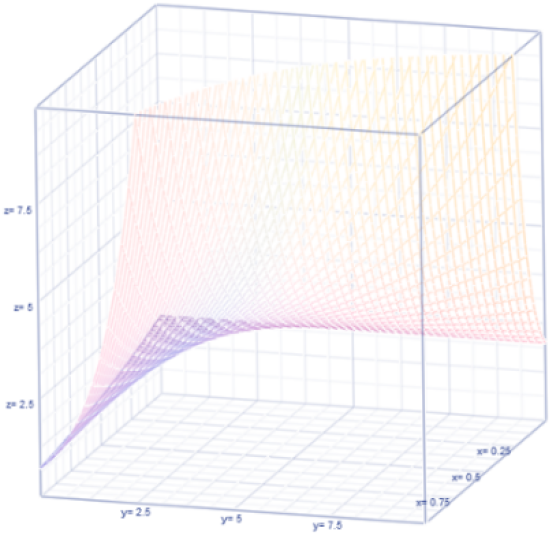
Odds for Asymptotic

Recall that 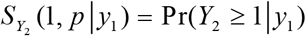 is the likelihood for the existence of asymptomatic presentation of COVID-19 in the ship. The hazard in that situation (that is, with *r* = 1) is

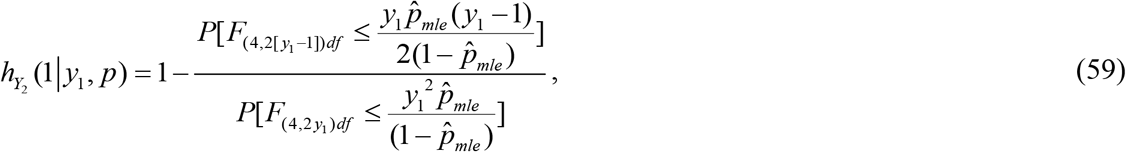

Where 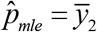. A popular statistical concept in the business world (Khokhlov, 2016 for details), *Tail Value at Risk* (Tar) is

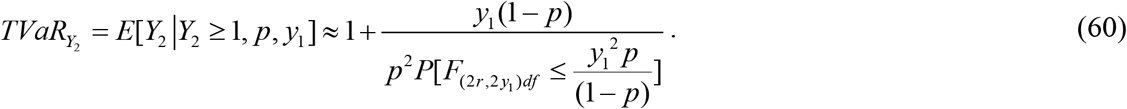

Similarly, all the Bayesian results for the binomial random variable, *y*_3_ are easily derivable by interchanging*γ* and*δ* in all above expressions. The survival function of the random number, *Y*_3_ with symptomatic symptoms is

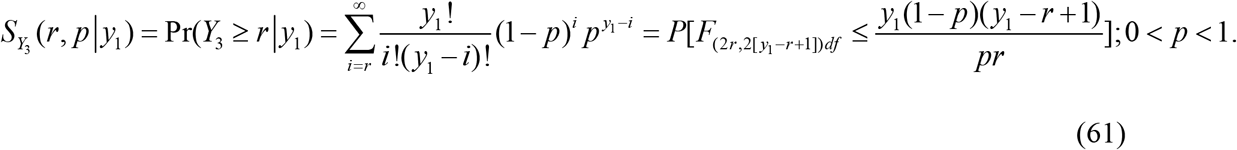

The hazard rate, *h*(*y*)for the symptomatic sign is

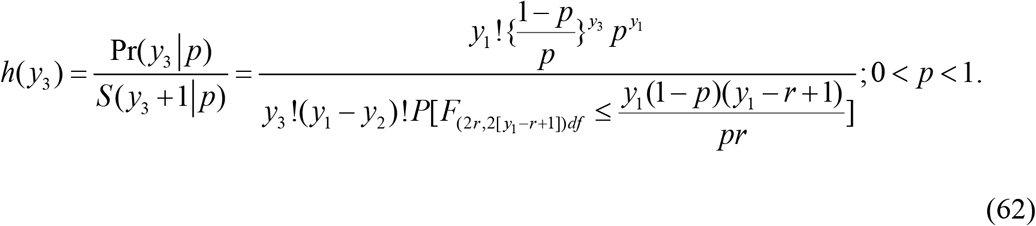

The binomial distribution of those with symptomatic signs has a finite *memory*

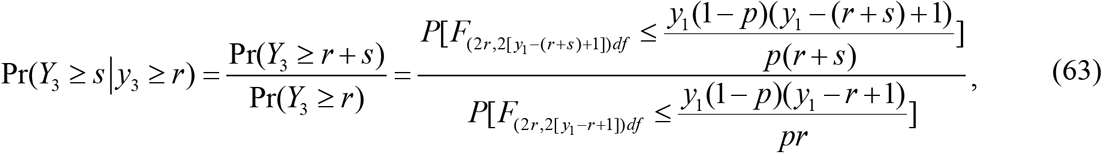

confirming that the usual binomial probability trend of those with symptomatic signs does possess a finite memory. The conditional odds, for a fixed *y*_1_, for *safe* symptomatic symptom are

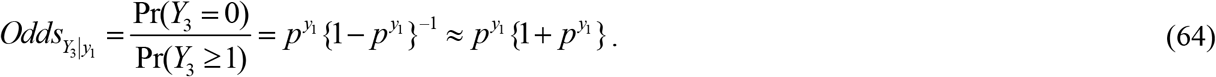

The unconditional odds for safe symptomatic symptom are

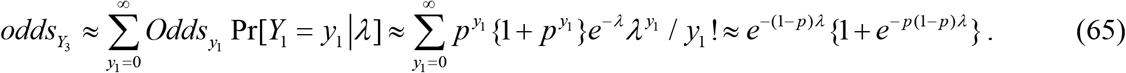

A comparison of 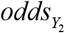 and 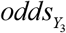 suggests the odds ratio,

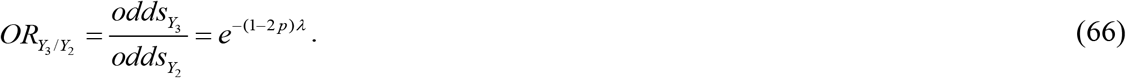

See Figure 5 for the configuration of the isomorphic factor, *e*^−(1−2 *p*)*λ*^.

**Figure 5.**
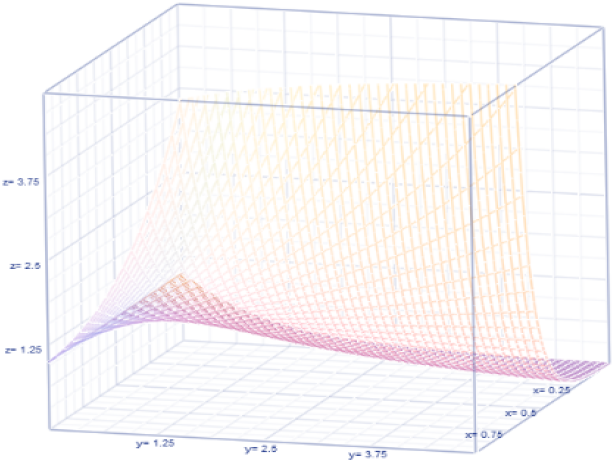
The configuration isomorphic factor *e*^−(1−2 *p*)*λ*^

Recall that 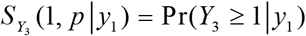 is the chance for the existence of symptomatic symptom of COVID-19. The hazard in that situation (that is, with *r* = 1) is

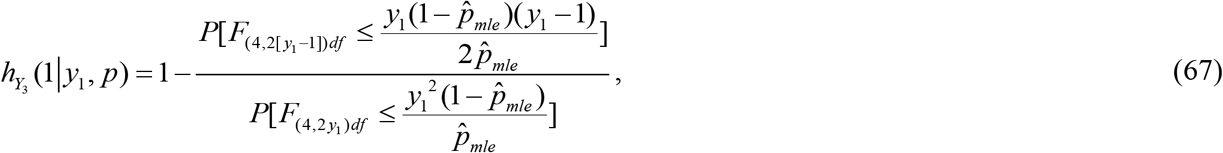

where 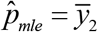. The *Tail Value at Risk* (TVaR) is

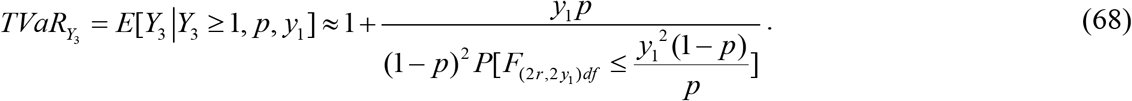

